# Sequence-based chromatin activity modeling and regulatory impact prediction of genetic variants in farmed animals using deep learning

**DOI:** 10.1101/2025.02.21.639224

**Authors:** Dat Thanh Nguyen, Tim Martin Knutsen, Simen R. Sandve, Sigbjørn Lien, Lars Grønvold

## Abstract

Non-coding genomic variations are crucial for the genetic regulation of traits; however, their functional impact in farmed animals remains underexplored due to limited genomic resources and the absence of tailored computational tools. Here, we present a deep learning-based framework that utilizes functional genomics data to generate genome-wide predictions of the regulatory impact of non-coding variants in cattle, chicken, pig, and Atlantic salmon. By leveraging chromatin profiles such as ATAC, DHS, and ChIP-seq data, we train and optimize separate deep networks for each species, achieving robust sequence modeling accuracy specific to each. Motif analysis confirms that the models capture regulatory grammar, while in silico saturation mutagenesis experiments provide meaningful interpretations of the functional impact of putative causal variants. Furthermore, functional scores derived from these models predict eQTL causal variants and enhance genomic prediction performance. Our findings highlight the transformative potential of sequence to function models in prioritizing causal variants and improving genomic prediction for livestock and aquaculture.

## 1 Introduction

Non-coding genomic variations are known to constitute the majority of disease- and complex trait-associated single nucleotide polymorphisms (SNP)^[1,2,3]^. Despite the success in identifying these variants, it remains challenging to accurately determine which ones are causal based solely on association results, as many neutral genomic variants are also significantly associated with traits in GWAS due to linkage disequilibrium (LD)^[4]^. To complement the limitations of population-based association studies, high-throughput functional assays of regulatory elements such as DNase I hypersensitive sites (DHS)^[5]^, transposase-accessible chromatin with sequencing (ATAC-seq)^[6]^, and chromatin immunoprecipitation sequencing (ChIP-seq)^[7]^ have become invaluable tools for detecting cis-regulatory elements (CRE) and prioritizing putative non-coding causal variants.

The advancement of deep learning, combined with the availability of large-scale functional annotations from projects like ENCODE^[8]^ and Roadmap Epigenomics^[9]^, has transformed the training of deep models for predicting chromatin states and assessing the impact of non-coding variants from DNA sequences in human genomics. In principle, deep neural networks are initially trained to differentiate putative regulatory sequences from background DNA sequences^[10,11,12]^, or to directly predict experimental read coverage from high-throughput functional assays^[13,14]^. Subsequently, these models are applied to predict the regulatory impact of any genetic variants by analyzing the disparities between reference and alternative alleles. Due to rapid advancements in artificial intelligence, deep models for regulatory genomics have evolved into various architectures that continuously improve modeling performance. Notable architectures include DeepSEA and Bas-set, which are purely convolutional neural networks (CNNs)^[10,12]^; Basenji, which employs dilated CNNs^[13]^; DanQ and DeepATT, which are hybrid CNNs with re-current neural networks (RNNs)^[11,15]^; and DeepFormer and Enformer, which are hybrid CNNs with transformers^[14,16]^.

In the realm of livestock and aquaculture genomics research, understanding the functional impact of non-coding variants is equally important. Variants in cis-regulatory elements can significantly influence transcriptional regulation, affecting traits of economic importance^[17]^. However, the exploration of functional non-coding variants in these species has fallen behind human genomics, primarily due to the absence of specialized functional genomics resources and tailored computational tools.

Genomic prediction has revolutionized livestock breeding by enabling the selection of individuals based on dense genomic marker information, thus increasing the precision of breeding value predictions for economically important traits^[18,19,20,21]^. Despite its transformative success, genomic prediction primarily relies on SNPs as markers without considering the functional impact of the variants which potentially help to improve genomic prediction performance. The reason for this stem from the lack of effective methods for prioritizing functional variants, highlighting the urgent need for approaches to predict the impact of regulatory sequence variation in farmed animals.

In this work, we develop deep learning-based sequence models that utilize large-scale functional genomic datasets from multiple species, including cattle, pig, chickens, and Atlantic salmon. These datasets feature ATAC, DHS, histone modification, and transcription factor ChIP-seq profiles^[22,23]^ to prioritize functional variants. We evaluate variant effect predictions using expression quantitative trait loci (eQTL) data from large consortium projects, demonstrating that the models effectively predict putative functional variants. The predicted scores further enhances genomic prediction as illustrated in a case study with Atlantic salmon. Overall, our proposed functional scores show significant potential for prioritizing causal variants and enhancing genomic prediction practices in livestock and aquaculture breeding.

## 2 Materials and Methods

An overview of the study is shown in Figure 1, which comprises three main steps: (i) data collection and preprocessing, (ii) deep learning model training and optimization, and (iii) regulatory impact inference. First, chromatin profile peaks from multiple farmed animal species are collected and preprocessed to obtain sequences and labels for deep model training (Figure 1 A, B). Next, we evaluate the performance of two widely used deep learning architectures for regulatory sequence modeling and variant impact prediction, DeepSEA^[10]^ and DanQ^[11]^, by testing different learning rates and applying the commonly used hold-out chromosome validation approach to identify the optimal model for each species (Figure 1 C). Finally, the best-performing models are used to infer the impact of regulatory variants for the corresponding species (Figure 1 D). Further details are provided in the following sections.

**Figure 1:**
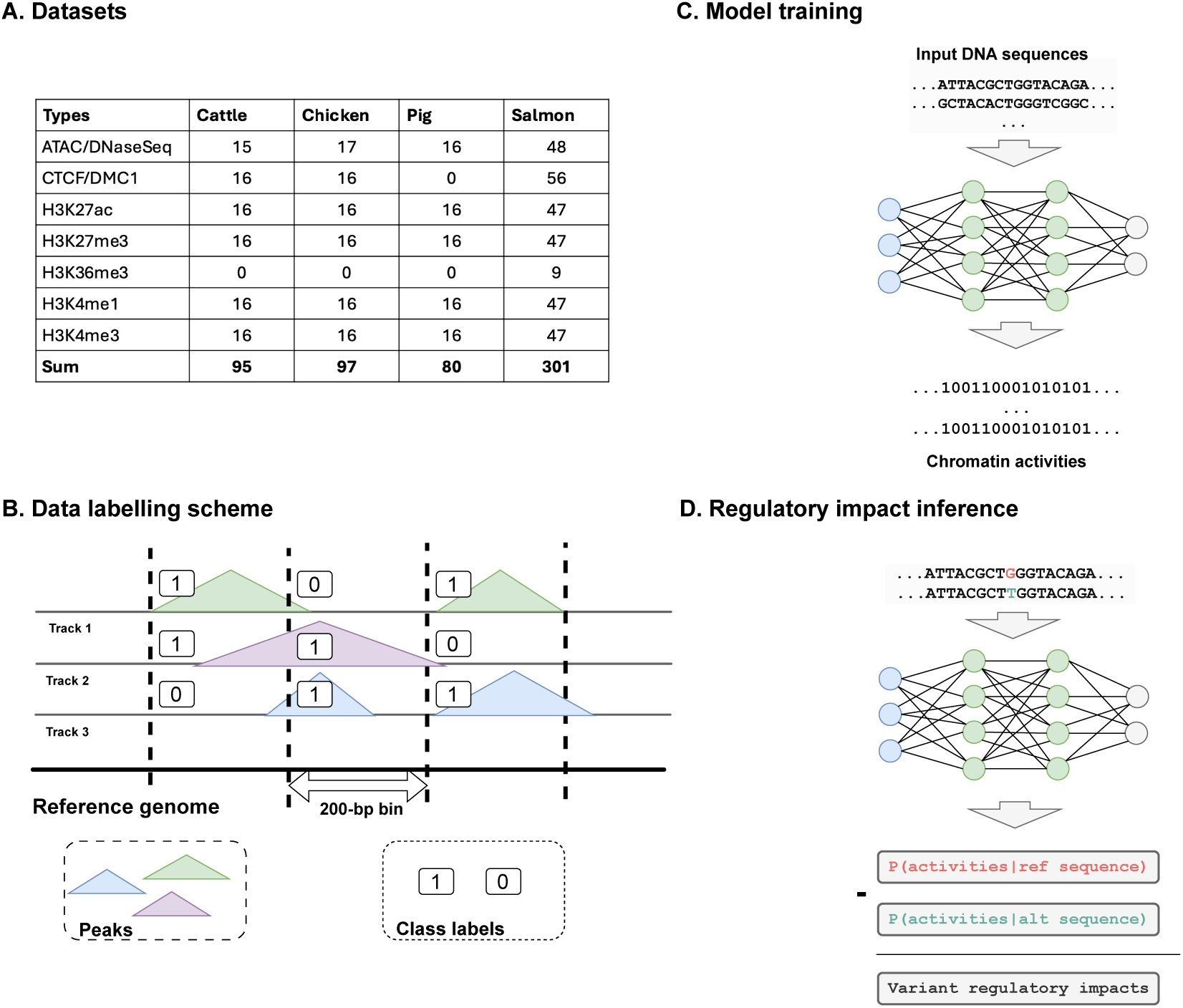
Overview of the study. A) Details of chromatin profiles used for each species B) Data preprocessing scheme to obtain sequence labels C) Deep model training and optimization and D) Variant impact inference scheme.

### 2.1 Training data

To train the machine learning models, we gather chromatin feature peaks from cattle (Bos taurus), pig (Sus scrofa), chicken (Gallus gallus), and Atlantic salmon (Salmo salar), sourced from the FAANG^[22]^ and AQUA-FAANG^[23]^ projects. For cattle, pig, and chicken, we utilize 95, 80, and 97 chromatin profiles, respectively, including data on histone modifications, CTCF ChIP-seq, DHS, and ATAC-seq across eight tissues: liver, lung, spleen, skeletal muscle, subcutaneous adipose, cerebellum, brain cortex, and hypothalamus. For Atlantic salmon, we use 301 profiles from histone modification, ChIP-seq, and ATAC-seq experiments across six tissues: brain, liver, gill, gonad, and muscle. Detailed statistics are provided in Figure 1 A.

For each species, we start by downloading the appropriate genome assembly version used for peak calling from the Ensembl database and then divide the genome into 200-bp bins. Specifically, we utilize the ARS-UCD1.2 genome for cattle, Sscrofa11.1 for pig, GalGal6 for chicken, and Ssal v3.1 for Atlantic salmon. DNA sequences are extracted from each bin and encoded using a one-hot scheme: A = [1,0,0,0], C = [0,1,0,0], G = [0,0,1,0], T = [0,0,0,1], and N = [0,0,0,0]. To meet the input context length required by DeepSEA and DanQ, we include the sequences of two bins upstream and downstream of each target bin, concatenating them to create a final 1000-bp training sequence. To label each DNA bin, we use BEDTools^[24]^ to evaluate overlaps between bin coordinates and chromatin peak regions. Following the DeepSEA approach^[10]^, a bin is assigned a label of 1 if at least 50% of its length overlaps with a peak region; otherwise, it is assigned 0. This process generates a label vector for each 200-bp bin, with the vector size corresponding to the number of chromatin profiles, as illustrated in Figure 1 B.

All autosomes, excluding those designated for validation and testing, are used for training. Specifically, chromosome 21 is reserved for validation and chromosome 25 for testing in cattle, chicken, and salmon, while chromosomes 16 and 17 are reserved for validation and testing, respectively, in pig.

### 2.2 Model architectures

**DeepSEA**^[10]^ utilizes a three-block convolutional architecture for feature extraction. The first convolutional block consists of 320 filters with a kernel size of 8, followed by ReLU activation, max pooling with a window size of 4, and dropout with a probability of 0.2. The second block extends this architecture with 480 filters of the same size, applying identical pooling operations and maintaining the dropout rate of 0.2. The third block further increases the complexity by incorporating 960 filters, and includes a higher dropout rate of 0.5 to better address over fitting. Following the convolutional and pooling operations, the network flattens the output and processes it through two fully connected layers to produce the final classification outputs.

**DanQ**^[11]^ begins with a convolutional block that applies 320 filters of size 26 to the input sequences, followed by ReLU activation, max pooling with a window size of 13, and dropout with a probability of 0.2. The output of this block is then fed into a bidirectional LSTM (Long Short-Term Memory) layer, which has 320 hidden units and processes the sequence data to capture temporal dependencies. Dropout with a probability of 0.5 is applied to the LSTM output to prevent over fitting. The output is then flattened and passed through two fully connected layers for classification.

### 2.3 Model training and evaluation

Deep learning models are trained using TensorFlow v2.12.0 on an NVIDIA Quadro RTX 8000 GPU, with parameters randomly initialized using TensorFlow’s default settings. During optimization, Binary Cross-Entropy (BCE) loss function is used as the objective function defined as BCE = 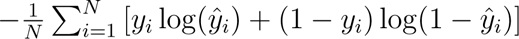, where *y_i_* is the true label, *ŷ_i_* is the pre-dicted probability, and *N* is the number of samples. The Adam optimizer^[25]^ is applied with a weight decay of 1 *×* 10*^−^*^6^ and a batch size of 1024. To identify the optimal learning rate for each method-dataset pair, we evaluate four candidate values: 1 *×* 10*^−^*^3^, 5 *×* 10*^−^*^4^, 1 *×* 10*^−^*^4^, and 5 *×* 10*^−^*^5^. Models are trained for up to 100 epochs, with early stopping applied if the validation loss does not improve for 5 consecutive epochs.

Finally, model performance is assessed on hold-out test sets. The Area Under the Receiver Operating Characteristic curve (AUROC) is used to select the best model for each species. Both forward and reverse DNA sequences are used during training, while only forward sequences are considered for validation and final evaluation.

### 2.4 Regulatory impact prediction

Regulatory impact prediction involves evaluating the potential effects of genetic variants on regulatory activities^[26,10,12]^. For each variant, two 1000-bp sequences centered on the variant position are extracted from the reference genome: one carrying the reference allele and the other carrying the alternative allele. These sequences are used as input to the trained model, which calculates the probabilities *P*_reference_ and *P*_alternative_ that the reference and alternative sequences belong to active chromatin regions, respectively. The regulatory impact of the variant is inferred as the difference between these probabilities: Δ*P* = *|P*_reference_ *− P*_alternative_*|*.

### 2.5 Fine-mapped eQTL variant classification

To assess the utility of the predicted functional scores, we adopt the methodology described by Avsec et al.^[14]^, focusing on putative eQTL causal variants. Specifically, we utilize the SuSiE^[27]^ fine-mapped results across multiple tissues provided by the PigGTEx consortium^[28]^. We include only tissues with at least 500 variants having a posterior inclusion probability (PIP) exceeding 0.9 in a credible causal set, resulting in 13 tissue-specific causal variant datasets. Negative sets are matched by sampling variants with PIP *<* 0.01 but *|Z*-score*| >* 4 for the same gene. If no such variants are available, alternatives are selected from genome-wide variants with PIP *<* 0.01 and *|Z*-score*| >* 6.

Using predicted functional scores, we then train separate random forest classifiers for each tissue to distinguish between positive and negative variant sets using ten-fold cross-validation. Default hyperparameters from scikit-learn are used, and 50 iterations of stochastic cross-validation and random forest fitting are performed to estimate model accuracy and its standard deviation.

### 2.6 Genomic selection analysis

As genomic prediction is a core application of livestock and aquaculture genomics, we further evaluates the potential of functional scores for prioritizing SNP sets within genotyping arrays. The Atlantic salmon SNP array, developed by AquaGen, comprises 70,000 markers (70k array) is downscaled to 9,073 SNPs, corresponding to a density of one SNP per 250 kb. Seven SNP sets are constructed: five randomly selected subsets from the 70k array, one functional set comprising the top 9,073 SNPs with the highest functional scores across the array, and one functional set consisting of the top 9,073 SNPs with the highest functional scores stratified by 250 kb windows (i.e., selecting the SNP with the highest functional score within each bin). Functional scores for ranking SNPs are computed as the average scores derived from epigenomic profiles of brain, gonad, and liver tissues of male Atlantic salmon.

Two widely utilized genomic prediction methods are applied including GCTA-Yang, a SNP-BLUP approach implemented in GCTA; and GCTB-Bayesian, a Bayesian framework implemented in GCTB. The GCTA-Yang method adopts a genomic best linear unbiased prediction (GBLUP) framework, assuming equal contributions of all SNPs to the genetic architecture of the trait and leveraging a genomic relationship matrix to predict genetic values^[29]^. Conversely, the GCTB-Bayesian method fits all genotyped markers simultaneously, accommodating heterogeneity in trait genetic architecture and providing a more flexible framework for genome-wide association studies and genomic prediction^[30]^.

The dataset used in this analysis comprises Atlantic salmon from three cohorts: AS19, AS20, and AS21. Each individual is genotyped for SNPs and assigned a binary phenotype representing late maturation. Covariates, such as year and age are included to account for environmental and batch effects. The models are trained using data from the combined AS19 and AS20 cohorts and validated on the genetically distinct AS21 cohort, ensuring a robust assessment of model generalizability.

Training is conducted across all SNP sets and prediction methods. Correlations between observed phenotypes and predicted phenotypes are subsequently calculated for each SNP set, providing a quantitative measure of the effectiveness of functional scores in enhancing genomic prediction.

## 3 Results

### 3.1 Optimizing chromatin activity modeling for farmed species

To optimize chromatin activity modeling, we evaluate two widely used deep learning architectures, DeepSEA and DanQ, across multiple learning rates. The results are summarized in Figure 2 and Table S.1. Overall, both architectures demonstrate high performance across species, achieving robust AUROC scores. Notably, DanQ dominantly outperforms DeepSEA with slightly higher AUROC scores across species and learning rates tested. For instance, in Cattle, the median AUROC score for DanQ ranges from 0.9066 to 0.9110, while DeepSEA achieves scores between 0.9010 and 0.9076. Similarly, in chicken, DanQ scores range from 0.8700 to 0.8825, compared to 0.8616 to 0.8800 for DeepSEA. The advantage of DanQ may be attributed to its ability to model the temporal dynamics of regulatory vocabularies^[11]^.

**Figure 2:**
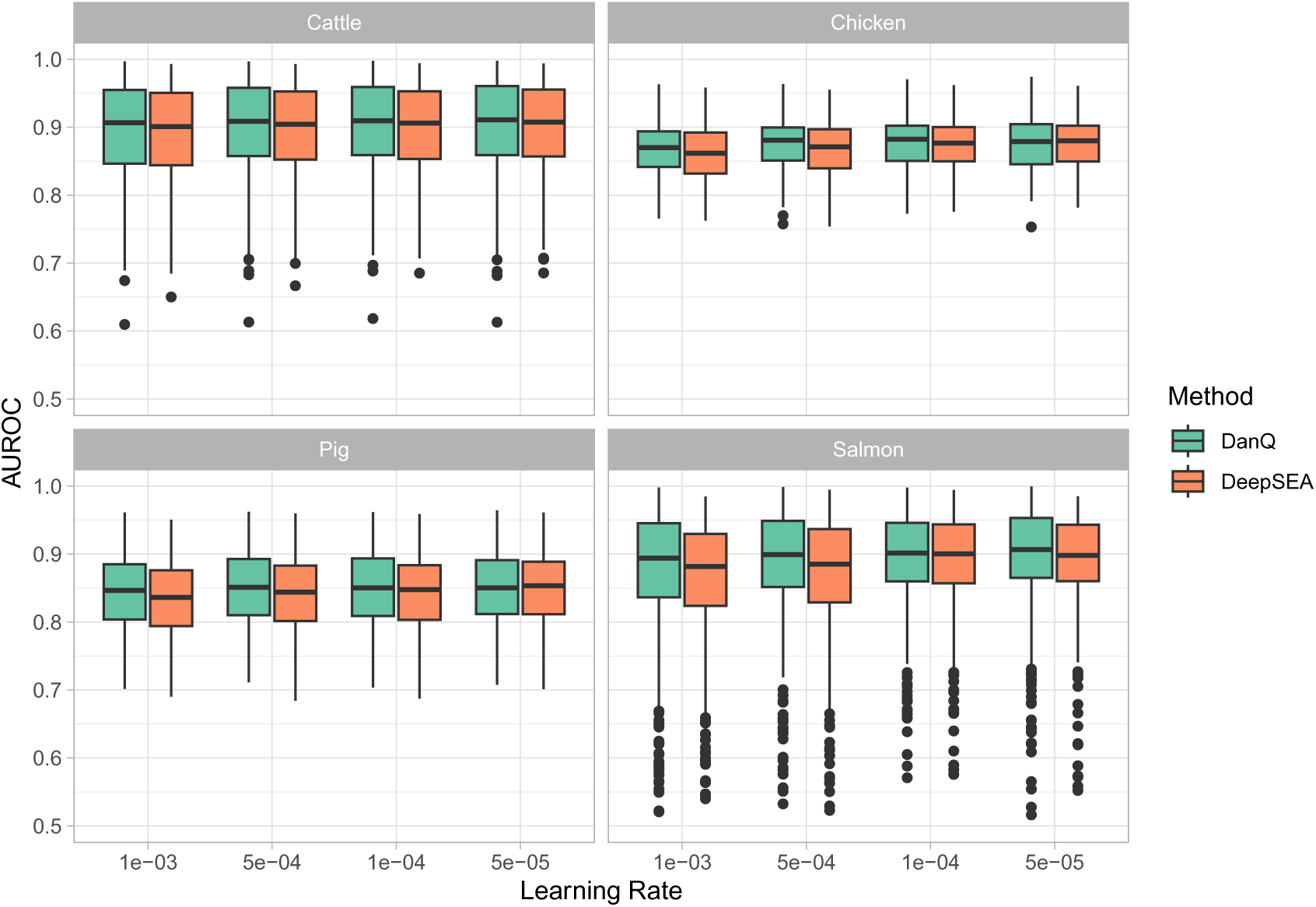
Performance comparison (AUROC scores) across learning rates of DeepSEA and DanQ in four species: cattle, chicken, pig, and salmon.

Building upon these findings, we choose DanQ as the primary method for evaluation by selecting the best-performing DanQ model for each species (Table S. 2). For cattle, the DanQ model trained with a learning rate of 5 *×* 10*^−^*^5^ achieves a median AUROC score of 0.9110. Similarly, in chicken, the best DanQ model, trained with a learning rate of 1 *×* 10*^−^*^4^, records a median AUROC score of 0.8825. For pig, the highest-performing DanQ model utilizes a learning rate of 5 *×* 10*^−^*^4^, achieving a median AUROC of 0.8512. Lastly, the salmon DanQ model trained at a learning rate of 5 *×* 10*^−^*^5^ achieves a median AUROC score of 0.9065.

To gain deeper insights, we explore the variability in performance across epigenetic profiles. As shown in Figure S.2, the AUROC scores exhibit significant variation depending on the data type and species. For instance, in cattle, H3K27ac and CTCF data achieve the highest median AUROC scores, while in salmon, H3K4me3 and ATAC/DNase stand out as the best-performing features. Chicken and pig display intermediate performance levels across most features. These results suggest that certain epigenetic features are more informative than others for regulatory activity modeling in the different species. Overall, these results indicate that we have achieved high accuracy, but the effectiveness of regulatory activity modeling varies across species and type of epigenetic features.

### 3.2 Deep models learn transcription factor binding motifs

To verify that the deep models effectively learn regulatory grammar, we conduct motif analysis. Following the approach outlined in the DeepBind method^[31]^, for each species, we convert the weight matrices from the first convolutional layer of the trained deep models into position-specific weight matrices (PSWMs). These learned motifs are then compared against the JASPAR 2022 CORE non-redundant v2 database using the TOMTOM algorithm^[32]^ implemented in the MEME Suite web service^[33]^.

Among the 320 motifs learned by each DanQ model, 49, 51, 63, and 34 motifs significantly match known motifs in the target database (E-value ¡ 0.01) for cattle, chicken, pig, and salmon, respectively. Among the significant matched motifs in cattle, we select four representative motifs and visualize them for illustration, as shown in Figure 3. These findings suggest that the trained models effectively learn regulatory vocabulary. However, the number of matched motifs is substantially lower than the 166 out of 320 matches reported for the human DanQ model^[11]^. This disparity may reflect the under-representation of farmed animal motifs in current databases and the limitation in term of training data for these species.

**Figure 3:**
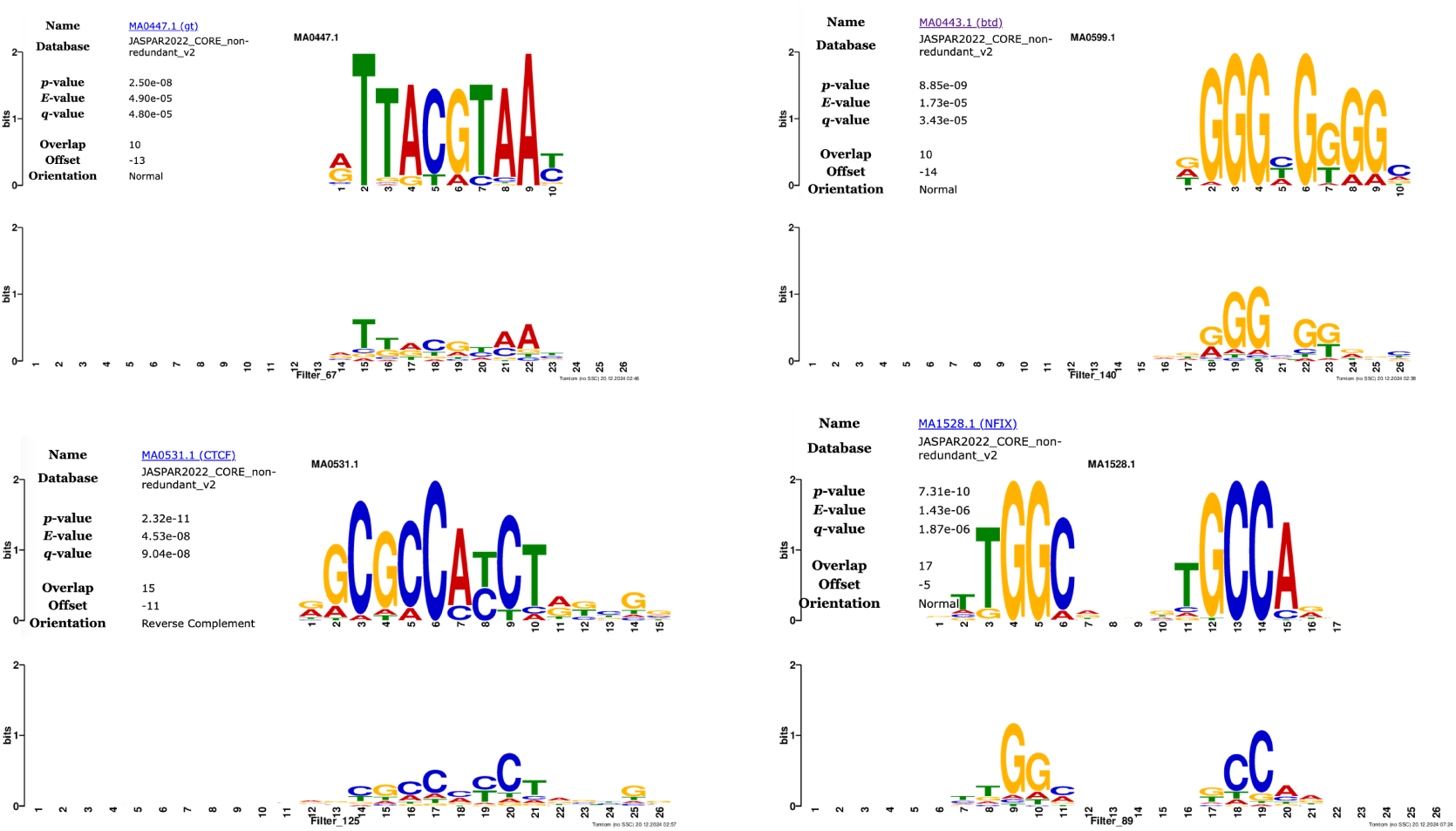
Visualization of four motifs GT, BTD, CTCF, and NFIX in cattle.

### 3.3 Functional scores predict eQTL causal variants

The utility of the predicted functional scores is evaluated through random forest classification across 13 pig tissue-specific datasets, distinguishing putative eQTL causal variants from matched negative variants. The mean AUROC scores from 50 iterations of ten-fold cross-validation vary across tissues, highlighting differences in predictive performance. The AUROC scores range from 0.7469 in spleen, which demonstrates the strongest predictive ability, to 0.5226 in uterus, indicating lowest performance. Other tissues exhibit intermediate levels of predictive accuracy, with scores of 0.6018 (lung), 0.5541 (testis), 0.5583 (adipose), 0.5707 (embryo), 0.5519 (ileum), 0.5895 (ovary), 0.5572 (small intestine), 0.5611 (brain), 0.5526 (muscle), 0.5508 (liver), and 0.5261 (blood).

The overall mean AUROC score across all tissues is 0.5726, indicating moderate performance of the functional scores in predicting putative eQTL causal variants. The results, including the mean and standard deviation of model performance, are detailed in Figure 4. These findings suggest that the predicted functional scores are correlated with eQTL predicted causal variants, although the strength of correlation varies across tissues.

**Figure 4:**
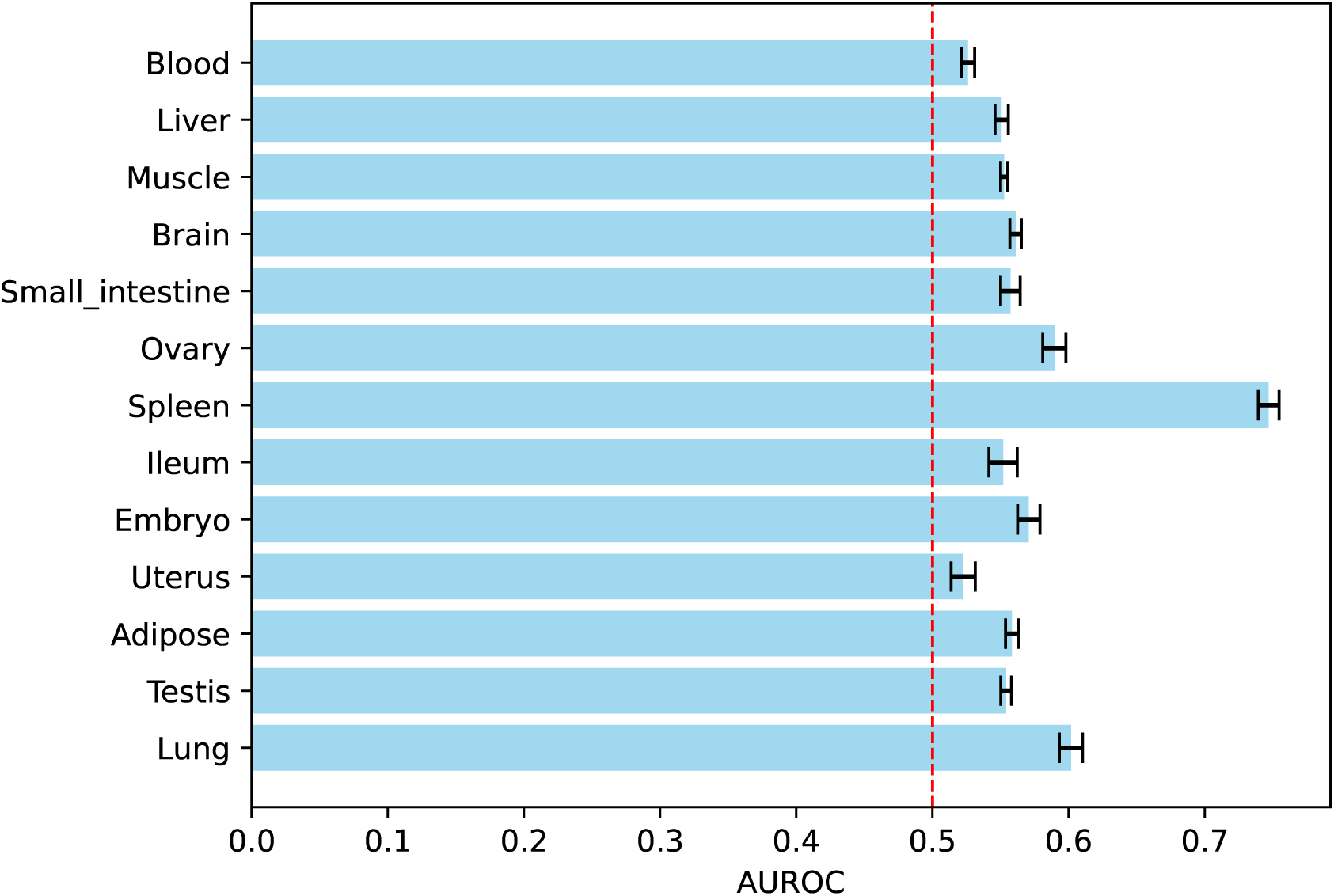
Mean and standard deviation of AUROC scores for 50 random forest eQTL causal variant classifiers across 13 pig eQTL datasets.

### 3.4 In silico saturation mutagenesis aids functional interpretation of putative causal variants

A trained model can be used to predict the functional activity of any given sequence, providing a powerful method for understanding and utilizing the regulatory patterns it has learned. In silico saturation mutagenesis experiments, which involve testing every possible mutation in a sequence, serve as an effective tool for identifying the specific nucleotides responsible for functional activity^[12]^. This approach is similar to standard functional impact prediction, where the difference in probabilities between two genotypes is computed. However, the inference is not limited to the variant position but extends to all possible variants within the targeted region.

We apply this strategy to investigate the putative regulatory variant rs133257289 in cattle, which is the top colocalized SNP of the cis-eQTL of DGAT1 and the GWAS association with protein yield^[34,35]^. In Figure 5, we present heat maps showing the change in predicted chromatin accessibility (mean of liver ATAC-seq tracks) due to mutations at each position, replacing the original nucleotide with each alternative for sequences surrounding the variant. These maps highlight the nucleotides most crucial to a sequence’s activity. For each position, we assign two scores: the loss score, which measures the largest possible decrease in activity, and the gain score, which measures the largest increase.

**Figure 5:**
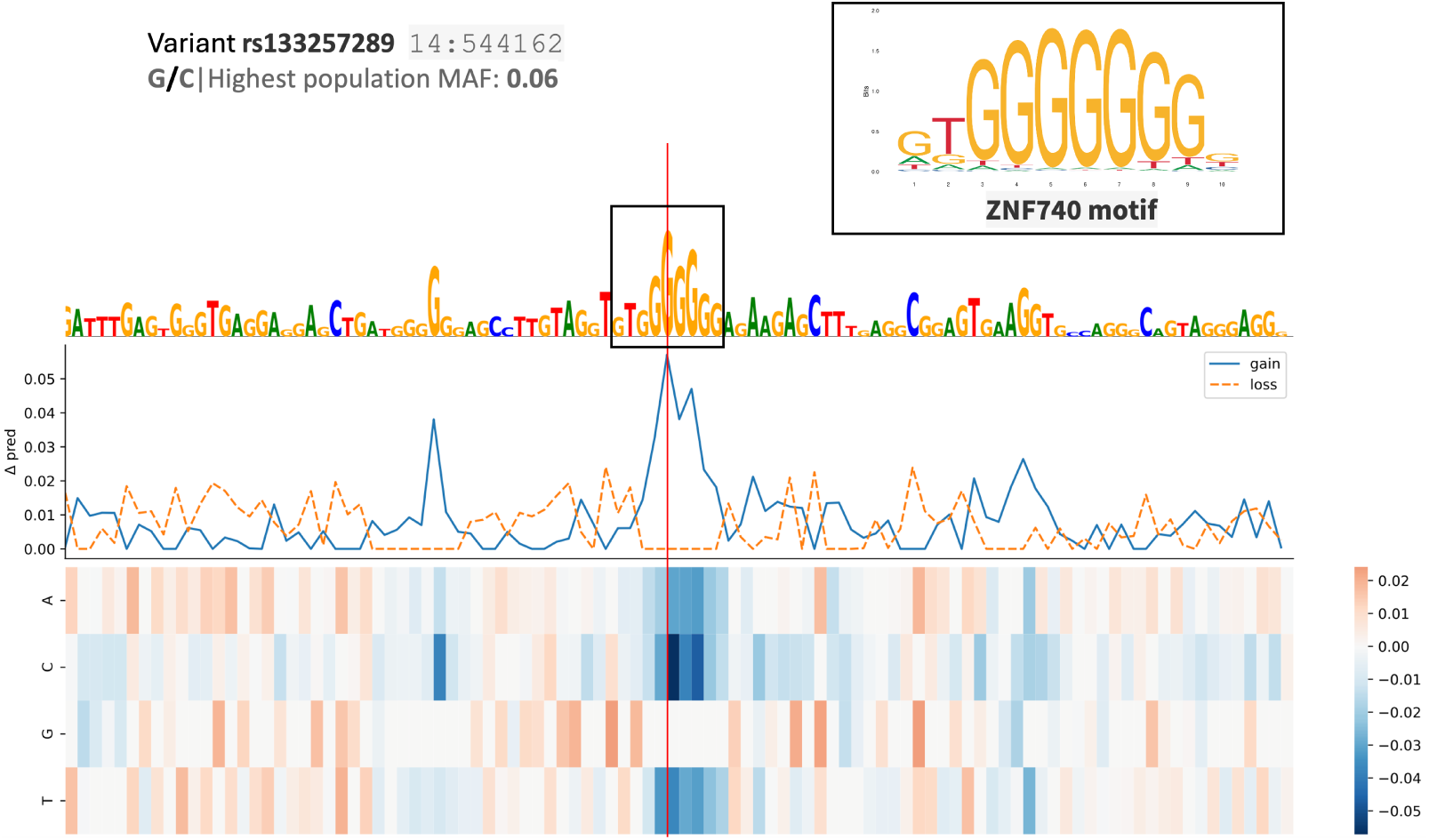
In silico saturation mutagenesis experiments of rs133257289, a putative regulatory variant in cattle, which is the top colocalized SNP for the cis-eQTL of DGAT1 and the GWAS association with protein yield.

High gain scores correspond to positions associated with the ZNF740 motif, where mutations disrupt the motif and increase chromatin accessibility. Notably, a G to C mutation aligns with the observed effect size of increased gene expression in the corresponding eQTL test in the liver ( p-value ¡ 1.2 *×* 10*^−^*^21^, effect size = 0.3123)^[34]^. These findings highlight the potential of in silico saturation mutagen-esis experiments using trained models for interpreting putative causal variants.

### 3.5 Functional impact scores improve genomic selection

To assess the impact of SNP selection strategies on genomic prediction accuracy, we evaluate the predictive correlation between observed and predicted phenotypes across various SNP sets and genomic prediction methods (Figure 6). The SNP sets include the full 70k SNP array, the top 9,073 SNPs selected based on functional scores (both genome-wide and within 250 kb bins), and random subsets of 9,073 SNPs derived from the 70k array.

**Figure 6:**
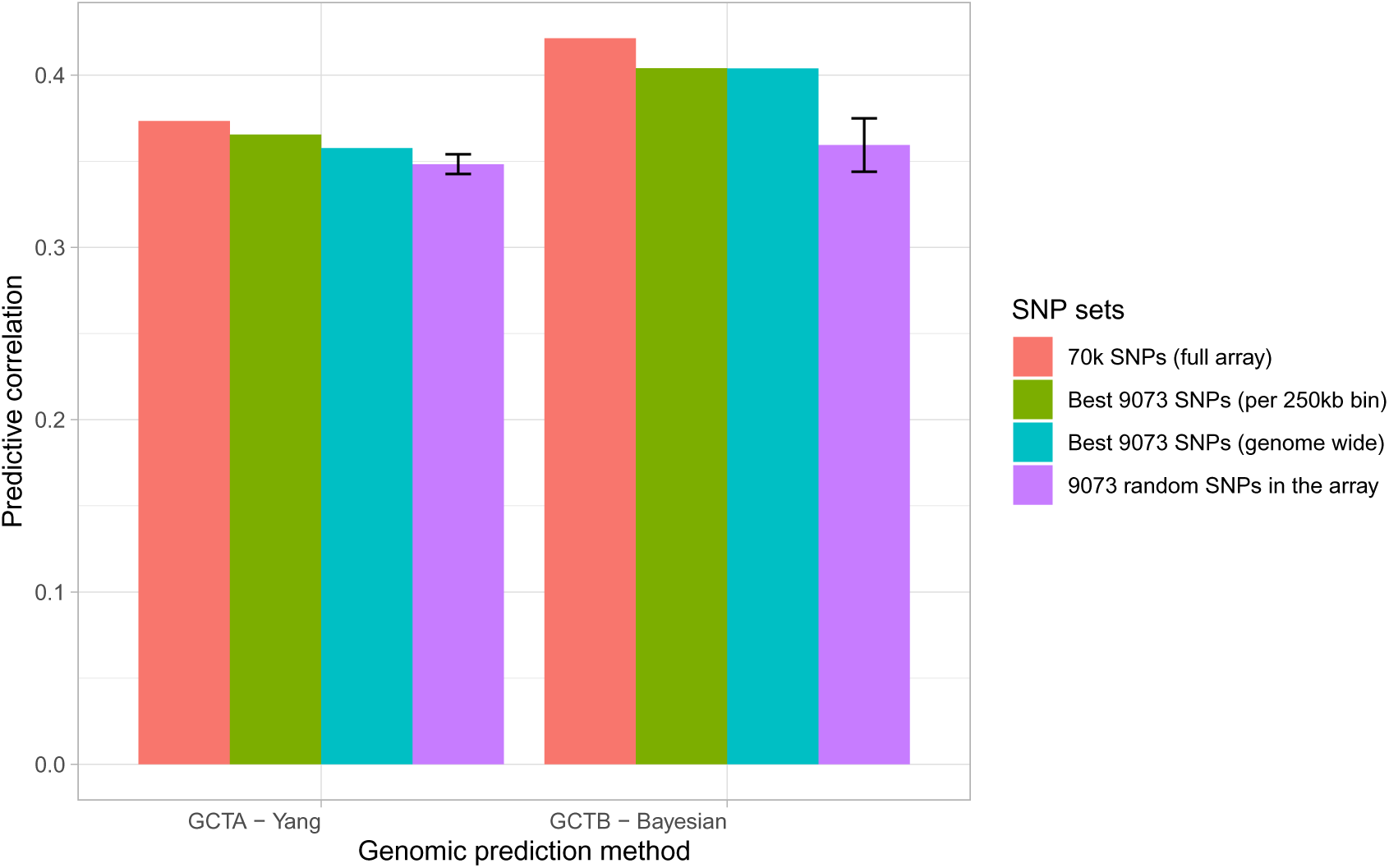
Genomic prediction performance of various SNP sets.

For the GCTA-Yang method, which assumes equal SNP contributions, the full 70k SNP array exhibits the highest predictive performance (correlation = 0.373), followed closely by the functional SNP set stratified by 250 kb bins (correlation = 0.365). SNP sets based on genome-wide functional scores (correlation 0.358) and random subsets of the array (correlation = 0.348 ± 0.0057) show slightly lower predictive correlations. In contrast, the GCTB-Bayesian method demonstrates improved predictive accuracy compared to GCTA-Yang across all SNP sets. The 70k SNP array achieves the highest predictive correlation (mean = 0.421), with functional SNP sets (both genome-wide and per 250 kb bin) showing comparable predictive performance (correlation = 0.404 for both SNP sets). Random SNP subsets yield the lowest predictive correlation under this method (correlation = 0.359 ± 0.0155).

Overall, these findings highlight the potential utility of functional scores in prioritizing SNPs for genomic prediction. Specifically, SNP sets derived from functional annotations perform on par with the full array, while significantly outperforming random SNP subsets. The results also demonstrate the advantages of the GCTB-Bayesian method over the GCTA-Yang approach, particularly in scenarios involving the maturation trait in Atlantic salmon.

## 4 Discussion

This study presents four deep learning sequence-to-function models, each trained on functional genomics data from cattle, pigs, chicken or salmon. To our knowledge this is the first time such models have been applied to farmed animals. We demonstrate the model’s ability to learn regulatory motifs and predict regulatory impact of non-coding variants. By enabling more precise identification of functional variants, these models can support more informed breeding decisions, ultimately contributing to sustainable improvements in livestock productivity and resilience. This work sets a foundation for further research and application of deep learning in the field of animal genomics.

We tried both the DeepSEA and DanQ deep learning architectures, settling on the DanQ architecture as it consistently returned higher AUROC scores in all species. This difference in performance between the architectures was also shown in the original DanQ model that was trained on human data^[11]^, indicating the finding generalizes well across species. The key difference between these architectures is the recurrent bLSTM block used in DanQ which has the ability to learn positional dependencies in the sequence, possibly reflecting the nature of the regulatory grammar.

Our motif analysis highlights the biological relevance of these models, demonstrating their ability to learn sequence patterns corresponding to known transcription factor binding sites. The DanQ model, with its single convolutional layer and long filters, is particularly suited for such analyses, as these filters effectively function as motif detectors. We identified motifs various significant motifs in all species, underscoring the models’ capability to capture regulatory elements even with limited training datasets in farmed animals. However, the reduced number of detected motifs compared to the human DanQ model may stem from key differences in the datasets. The original DanQ model was trained on a comprehensive functional data with large amount of TF ChIP-seq data making it to learn binding motifs specific to transcription factors in the dataset. In contrast, our models rely on broader chromatin accessibility profiles, which may limit motif specificity. Another factor to consider is the evolutionary divergence between species transcription factors in distant species compared to the reference database, such as salmon, may recognize motifs distinct from those in human or mouse, further influencing motif detection. These findings highlight the need for more comprehensive functional annotations in non-human species, which could enhance both motif discovery and overall model performance.

The application of in silico saturation mutagenesis demonstrates the practical utility of these models in interpreting putative causal regulatory variants. For instance, the analysis of rs133257289 in cattle reveals specific nucleotide changes that influence chromatin accessibility and align with observed eQTL effects, underscoring the potential of deep learning-based approaches for functional variant interpretation. Such methods provide invaluable insights for identifying candidate variants linked to economically important traits, paving the way for more targeted breeding strategies.

Moreover, our evaluation of functional scores in predicting eQTL causal variants in pig reveals moderate to high accuracy, with performance varying across tissues. Importantly, these functional scores demonstrate significant potential for enhancing genomic selection, as evidenced by their comparable performance to full SNP arrays and their superiority over randomly selected subsets as demonstrated in salmon. This finding suggests that functional annotations can be used to prioritize SNPs for functional-aware array design^[36]^, thereby improving the efficiency and accuracy of genomic prediction.

## 5 Conclusion

Overall, this study demonstrates the feasibility and utility of deep learning-based approaches for predicting the regulatory impact of non-coding variants in farmed species. By leveraging functional genomic data, our models achieve high predictive accuracy, uncover regulatory grammar, and provide actionable insights for genomic selection. The integration of functional annotations into breeding programs offers a promising approach for enhancing the precision and efficiency of livestock and aquaculture genomics. Future work should focus on expanding functional genomics resources for farmed species and exploring the application of these methods to other economically important organisms.

## Availability of data and materials

The deep learning models are freely available at https://github.com/datngu/DeepFARM. Data supporting the findings and source codes for data analyses and generating figures for this study are available at https://github.com/datngu/DeepFARM_paper. Functional datasets are available in its original publications^[22,23]^ at http://farm.cse.ucdavis.edu, and https://salmobase.org/. Cattle GTEX and Pig GTEX data are available at https://cgtex.roslin.ed.ac.uk/, and https://piggtex.ipiginc.com/. The datasets used in genomic prediction analysis are property of Aquagen AS that need permission to access.

## Authors’ contributions

DTN: conceptualized and implemented the methods; designed and analyzed the data; and wrote the manuscript with inputs from other co-authors. TMK and SL: conceptualized and implemented the genomic prediction experiment. LG, SRS and SL: contributed to discussions, manuscript review, and project administration. All authors read and approved the manuscript.

## Use of AI Software

Large language models were used to improve the wording and grammar of some texts, but not to generate new content.

## Acknowledgments

The authors would like to thank the AQUA-FAANG consortium for granting access to the dataset, the Orion HPC for providing computational resources. DTN also gratefully acknowledges the financial support from internal funding scheme at Norwegian University of Life Sciences (project number 1211130114), which financed the international stay at Queensland Institute of Medical Research, Brisbane, Aus-tralia.

