## Supplementary material for "Sequence-based chromatin activity modeling and regulatory impact prediction of genetic variants in farmed animals using deep learning": tables

### Supplementary tables

*Table S. 1 Median AUROC of DeepSEA and DanQ with various learning rates*

| LR | Spec | DanQ | DeepSEA |
|---|---|---|---|
| 1.00E-03 | Cattle | 0.9065707752 | 0.900988193 |
| 5.00E-04 | Cattle | 0.9085886943 | 0.9043951749 |
| 1.00E-04 | Cattle | 0.909432918 | 0.9061286703 |
| 5.00E-05 | Cattle | 0.9109503239 | 0.9076344934 |
| 1.00E-03 | Chicken | 0.8700395098 | 0.8615982768 |
| 5.00E-04 | Chicken | 0.8808054855 | 0.8711202143 |
| 1.00E-04 | Chicken | 0.8825036037 | 0.8765192327 |
| 5.00E-05 | Chicken | 0.8788894999 | 0.8800442025 |
| 1.00E-03 | Pig | 0.846664582 | 0.8361436374 |
| 5.00E-04 | Pig | 0.8512331557 | 0.844108463 |
| 1.00E-04 | Pig | 0.8502707891 | 0.8477812439 |
| 5.00E-05 | Pig | 0.8502366465 | 0.8534089627 |
| 1.00E-03 | Salmon | 0.8933912289 | 0.8816603276 |
| 5.00E-04 | Salmon | 0.897149215 | 0.8849771018 |
| 1.00E-04 | Salmon | 0.9010306001 | 0.9001452916 |
| 5.00E-05 | Salmon | 0.9065477151 | 0.8981181185 |

*Table S. 2 Best DanQ model for each species*

| LR | Method | Spec | Median_AUROC |
|---|---|---|---|
| 5.00E-05 | DanQ | Cattle | 0.9109503239 |
| 1.00E-04 | DanQ | Chicken | 0.8825036037 |
| 5.00E-04 | DanQ | Pig | 0.8512331557 |
| 5.00E-05 | DanQ | Salmon | 0.9065477151 |

*Table S. 3 Genomic prediction performance of GCTA Yang and GCTB Bayesian of various SNP sets*

| Dataset | Method | Score |
|---|---|---|
| All_SNPs | GCTA_Yang | 0.373394 |
| All_SNPs | GCTB_Bayesian | 0.421329 |
| Best_Score_Marker_per_250kb | GCTA_Yang | 0.365494 |
| Best_Score_Marker_per_250kb | GCTB_Bayesian | 0.403982 |
| Random_Affx_ID | GCTA_Yang | 0.343595 |
| Random_Affx_ID | GCTB_Bayesian | 0.348106 |
| Random_Affx_ID | GCTA_Yang | 0.354141 |
| Random_Affx_ID | GCTB_Bayesian | 0.374718 |
| Random_Affx_ID | GCTA_Yang | 0.353952 |
| Random_Affx_ID | GCTB_Bayesian | 0.373811 |
| Random_Affx_ID | GCTA_Yang | 0.341793 |
| Random_Affx_ID | GCTB_Bayesian | 0.360787 |
| Random_Affx_ID | GCTA_Yang | 0.347841 |
| Random_Affx_ID | GCTB_Bayesian | 0.339757 |
| Top_9073_SNPs_Across_All | GCTA_Yang | 0.357623 |
| Top_9073_SNPs_Across_All | GCTB_Bayesian | 0.403881 |
