## Supplementary figures and images for "Sequence-based chromatin activity modeling and regulatory impact prediction of genetic variants in farmed animals using deep learning"

### figures

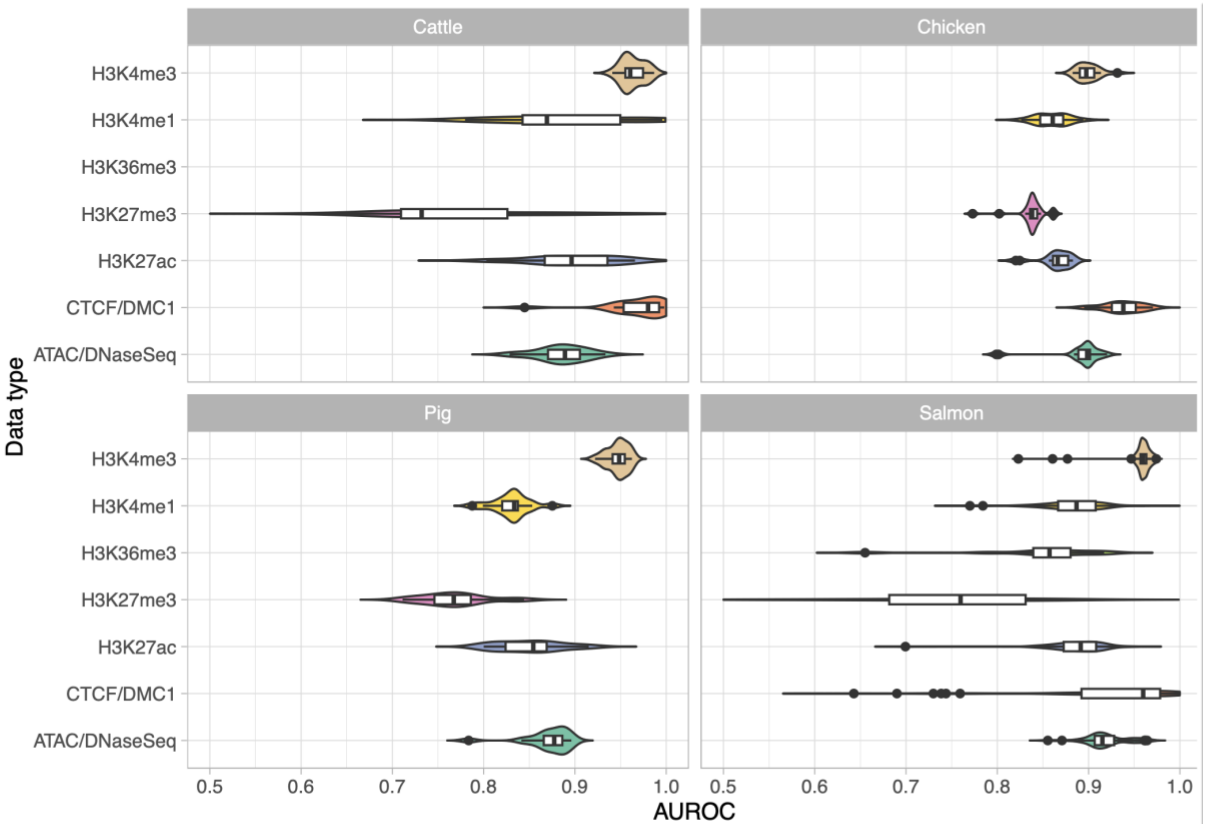


*Figure S. 1 AUROC performance across epigenetic profiles*
